# *Thalassolituus haligoni* BB40^T^, a model species for non-cyanobacterial diazotrophs within Oceanospirillales isolated from a Fjord-like Inlet in Kjipuktuk

**DOI:** 10.64898/2026.02.10.701148

**Authors:** Sonja. A. Rose, Stephanie. L.G. Duffy, Ludovic Pascal, Gwénaëlle Chaillou, Jennifer Tolman, Erin. M. Bertrand, Julie LaRoche

## Abstract

Marine non-cyanobacterial diazotrophs (NCDs) are recognized as globally distributed, however, few representatives have been isolated in pure cultures. *Thalassolituus haligoni* BB40^T^ was isolated from a fjord-like inlet within Kjipuktuk (Halifax), Nova Scotia. The genome displayed all necessary genes required for N_2_ fixation, and various carbon uptake pathways. The Gram-negative flagellated rod shape bacterium displayed significantly higher growth rates in medium amended with nitrate (NO_3_^-^) or ammonia (NH_3_), compared to dissolved N_2_, as the sole nitrogen source. Biological N_2_ fixation rates were detectable across all conditions, measuring a range from 9.34 × 10^-6^ to 1.4 × 10^-1^ fmol N cell^-1^ day^-1^. Growth of the isolate was successful between 4 °C up to 35 °C, with a T_opt_ of 20 °C for N_2_, and between 27 - 30 °C for fixed nitrogen (NO_3_^-^ and NH_3_). The closest relatives to *T. haligoni* BB40^T^, were found to be the uncultured Arc-gamma-03 (99% average nucleotide identity (ANI)) and *Oceanobacter antarcticus* wDCs (97% ANI). *T. haligoni* BB40^T^ also displays versatile capabilities for growth on various carbon, and nitrogen sources, and antibiotics. Collectively this study provides an in-depth physiological assessment of an Oceanospirillales diazotrophic species which we presently have limited knowledge of.

## Introduction

Nitrogen (N) serves as a main macronutrient within the marine life cycle, being incorporated into DNA, and proteins which are then used for growth across all aspects of the food web. Due to its importance, it is also rapidly taken up when available, making it a limiting nutrient across global oceans (1). Marine N_2_ fixation is a biological process which takes inert N_2_ gas and converts it into a fixed N form, ammonia (NH_3_), via the oxygen sensitive nitrogenase enzyme — providing fixed N input (2). This process is restricted to a select group of eukaryotes, bacteria and archaea, termed diazotrophs which can be further assigned into cyanobacterial and non-cyanobacterial diazotrophs (NCD)s (3–6). Taxonomic classification and detection of NCDs has been advanced by sequencing of the nitrogenase marker gene, *nifH* (7,8), clade-specific quantitative polymerase chain reaction (qPCR) assays (9–11), and more recently by assembly of metagenome assembled genomes (MAGS) (12) Recent studies (4,13–16), have shown NCDs to be globally distributed within marine systems, and to fix N_2_ in surface waters in the presence of fixed N, which is counterintuitive for the process given the energetic demands (17) (Mills et al., 2020). However, because few NCDs have been cultivated, their physiology and N_2_ fixation capacity remain poorly understood (4).

The family *Oceanospirillaceae* belongs to a diverse group of diazotrophic and non-diazotrophic species, with few cultured representatives. The family is comprised of rod-like flagellated heterotrophic Gram-negative bacteria that utilize a wide variety of carbon sources, ranging from C_4_ dicarboxylic acids to phenolic compounds and oil byproducts (18). To date, *Oceanospirillaceae* contains 17 genera, including *Oceanobacter, Oceanospirillum, Marinobacter* and *Thalassolituus* (18). Guanine and Cytosine (GC) content within *Oceanospirillaceae* ranges from 41 to 65 mol%, which contributes to the large diversity and various phenotypic characteristics seen across different genera.

The larger ubiquitous group of *Oceanobacter-*related taxa (ORT), consists of close phylogenomic relationships between *Oceanobacter, Thalassolituus* and *Oleibacter* (19). *Oceanobacter* sp. typically contain free-living flagellated motile cells (0.4 - 1.2 µm long and 2.0 - 4.0 µm wide), and are strict (micro-)aerobic halophiles (20). The genus tends to lack the *alkB* (alkane monooxygenase) gene, accumulate poly-β- hydroxybutyrate granules (PHB) and is represented by the non-diazotrophic species, *Oceanobacter kregii* (19,20). The genera *Thalassolituus,* typically contains free-living motile cells (1.2 - 2.5 mm long and 0.6 mm wide), are strict aerobic halophiles (18,21) and are represented by the non-diazotrophic species, *Thalassolituus oleivorans* (21). Given the complexities of classifications within the ORT, majority of taxonomy remains unresolved and heavily under debate for newly discovered taxa. Using genomics and physiology we introduce *Thalassolituus haligoni* BB40^T^, a marine heterotrophic Gram-negative facultative anaerobic NCD belonging to the *Oceanospirillales* order, which was isolated from an inlet along the Nova Scotian Coast (44°41′30″N, 63°38′30″W). Metabarcoding of the *nifH* gene indicated that the isolate is globally distributed, ranging from the Canadian Arctic Archipelago, down to the coastal and open ocean of the Eastern and Western Atlantic, to the Eastern Pacific, and South China sea (10). Data from the study below confirms that basal N_2_ fixation takes place in the presence of fixed N but is up to three orders of magnitude higher when grown without fixed N sources and in hypoxic conditions. This study provides a detailed physiological profile of the isolate, including growth rates under varying temperatures, N compounds and O_2_ concentrations. Phenotypic capabilities were verified using BIOLOG microarray plates across various C, N, and antibiotic substrates. Lastly, we provide a genomic pathway comparison of closely related species to *T. haligoni* highlighting the diversity of ORT NCD species. Collectively, the data presented herein proposes *T. haligoni* as a model species for *Oceanospirillacaea* NCDs.

## Materials and Methods

### Isolation and Initial growth

*Thalassolituus haligoni* BB40^T^ was isolated from a fjord-like inlet in Halifax, Nova Scotia, Canada, in January 2014 (44°41′30″N, 63°38′30″W; (10)). Water samples of 1, 5 and 10 and 60 m depth were aliquoted into 30 mL culture flasks and enriched with nutrients (*see* (10) for further isolation details). *nifH* PCR was used to monitor for diazotrophs, where positive cultures were single cell sorted using a BD influx single cell sorter and placed onto a N-free 1.2% agar plate (10). Visible colonies then underwent another plate transfer and were screened for *nifH*. Positive *nifH* colonies were transferred to YBC-II agar plates and supplemented with 15 mM sodium acetate until visible growth was seen on the plate (10). Culture samples underwent several rounds of genomic amplification (both short- and long read sequencing;(10)) to complete the genome and light microscopy samples were used to confirm the gram stain of the isolate (22).

### Genome sequencing, assembly and quality assessment

Short read sequencing for genome completion was extracted using the AllPrep DNA/RNA mini kit (QIAGEN), while DNA for long-read sequencing was extracted using QIAGEN’s Genomic-tops 20/G according to the manufacturers protocol ((10)). DNA concentrations were verified using the NanoDrop 2000c spectrophotometer (Thermo Fisher Scientific). Long-reads were further verified using agarose gel electrophoresis and final concentrations verified using the Qubit 4 fluorometer (Invitrogen; (10)). Genomic sequencing and library preparation was completed at the Integrated Microbiome Resource (IMR) at Dalhousie University using Illumnia MiSeq (300 – 300 bp paired end reads) and a Nanopore MinION Mk1 device (supplemental material 4 (10)). CheckM2 (v 1.0.2) was used to fully verify genome quality (supplemental material 4; (23)). Using a neural network, the genome was verified to be 99.77 %, with a contamination of 1.07. The average gene length was determined to be 327 bp, with 1 total contig, and a N50 of 4,376,545. The fully complete genome can be found under the accession number CP139482.1.

### Distribution and Comparison of *nifH*

*nifH* distribution comparison between *O. antarcticus* wDCs and *T. haligoni* BB40^T^ involved the same database search which was used in Rose et al., (2024) using the Qiime 2 V. 2019.7 (Martin et al., 2011; Comeau et al., 2017). Due to the similarities between the two *nifH* sequences (99% sequence ID) a pairwise identity of 100% was used for distinguishing between strains in distribution mapping. Presence was considered if raw reads were greater than 2 at the station of interest. All station information can be found in supplemental material 1.

### Physiological Characterization

#### BIOLOG

Phenotypic capabilities of *T. haligoni* BB40^T^ were tested using BIOLOG PM microarray Gram-negative plates (Plate numbers PM1, 2a, 3, 5-8, PM 19, 20), and standard aseptic growth techniques (Bochner 2009, Rose et al., 2024). Cultures for BIOLOG PM plates were initially grown using the same modified f/2 ASW 220 uM nitrate (NO_3_^-^), supplemented with 6.4 mM carbon as previous studies (99% purity; Fumarate, Thermo Scientific;(10)). Before plate inoculation, cell growth was tracked using the BD Accuri C6 autosampler flow cytometer (BD Biosciences) and stained with 1:1000 Sybr Green Dye II (Invitrogen). Cultures were then transferred to a modified minimal media reflective of the ASW ((24); supplemental material 2), however, void of either carbon or N sources, depending on the plate type being inoculated. Plate preparation followed standard protocol according to BIOLOG where cells were measured at OD590 nm using the Nanodrop Spectrophotometer 2000c in cuvette mode (Thermo Scientific). Once the desired density was reached, cells were transferred to the modified minimal media with either carbon (99% purity, 6.4 mM carbon, Fumarate) or fixed N (220 *µ*M NO_3_^-^; 99% purity; sodium nitrate, Sigma Aldrich) supplemented to allow for growth on the plate. Plates were then measured once a week using a SYNERGY H1 microplate reader (BioTek) at OD590 nm.

### Temperature range and Microscopy

The temperature range (4 − 40 °C) of BB40^T^ was examined under various N and oxygen conditions (N_2_, NO_3_^-^, NH_3_; hypoxic and oxic) to determine the cardinal temperatures (T_min_, T_opt_, T_max_) for the isolate. Cardinal temperatures within this study are defined as the lowest and highest temperature tolerance where growth is still detected (T_min_ and T_max_) and the optimal temperature (T_opt_) is the temperature at which highest growth rates were measured. Cells were grown in a modified f/2 artificial seawater (ASW) media supplemented with 6.4 mM carbon (99% purity Fumarate, Thermo Scientific) and 300 *μ*M phosphate with 1:1000 resazurin in 44 mL glass vials (*see supplemental of* (*10*) *for ASW recipe*). Cultures were stained with 1:100 Resazurin to track O_2_ concentrations via aerobic respiration, where blue indicates full oxygenation, pink indicates hypoxic and colorless indicates anoxia. Hypoxic conditions were created by flushing with N_2_ gas and sealing the vials with septum caps to remove all O_2_ from the flask until purple/pink in colour. After which, cells were added and a pink colour was reached and maintained within two hours of growth, signifying hypoxic conditions. Fixed N conditions used either 200 *µ*M NH_3_ (99% purity; Ammonium Chloride, Sigma-Aldrich), 220 *µ*M NO_3_^-^ (99% purity; sodium nitrate, Sigma Aldrich) or atmospheric N_2_. Hypoxic conditions included all three fixed N sources, while oxic conditions only included NH_3_ and NO_3_^-^ as the bacterium was unable to grow in oxic N_2_ conditions. Cultures were kept in exponential phase, and grown for a minimum of 9 generations, after which, cells were transferred to new culture tubes, with three replicates per temperature. Due to differences in growth rates, cell measurements were taken every 3 hours for fixed N and every 3 days for N_2_ cultures. Cell densities were recorded on the Novocyte flow cytometer and stained with 1:1000 Sybr Green Dye II (Invitrogen). Microscopy samples were collected, prepared and preserved in the exponential phase for all five conditions at 17 °C following protocols in (10). Microscopy samples were then examined at the Electron Microscopy Facility at Dalhousie University, Halifax, Nova Scotia, Canada.

Generation times were calculated using the slope of the curve on log scale, where N1 and N2 are initial and final cell densities (cell L^-1^) within the exponential phase of growth, and divided by the difference in time (Δt; days) and multiplied by 3.3 (Equation 1.1). Growth rates were then calculated by dividing 0.301 by the generation time, where 0.301 is log_10_2 (Equation 1.2):

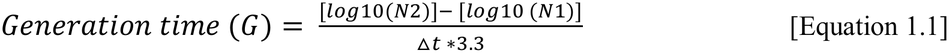

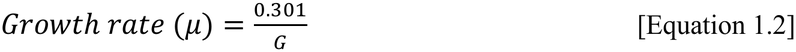

### Phylogenomic and Phylogenetic Analysis

Phylogenomic analysis was carried out using the published genome of *T. haligoni* BB40^T^ using 71 single-copy marker genes from Anvi’o v 7.8 using -- hmm-source- Bacteria_71 (*see methods* in (10,25)). Once best hits were established --return-best-hit, sequences were translated into amino acid sequences using --get-aa-sequences and the concatenated with --concatenate. The full list of single-copy core marker genes can be found in supplemental table 4. The phylogenomic tree was then curated using the concatenated protein fasta files using anvi-gen-phylogenomic-tree from the Anvi’o pipeline (25). The outgroup selected for both the phylogenomic, and phylogenetic tree was *Alcanivorax borkumensis* ABS183 due to it being a related member of *Oceanospirillales* but lies outside the ingroup of *Oceanospirillaceae*. The maximum likelihood phylogenomic tree was curated from the concatenated amino acid alignment using FastTree2 under the Jones Taylor-Thorton model, with approximation of among-site rate variation with 1,000 bootstraps. Labels for the phylogenomic tree were then modified in iTOL (26,27).

An average nucleotide identity (ANI) was conducted using Anvi’o v7.8 (25). External genomes were downloaded from bacterial and viral bioinformatics resource center (BV-BRC) and an external contigs database created using the external_contigsdb_anvio.sh script. Once populated, annotations were curated for each contig database, COG, KEGG, KOFam, and pfam databases (28–30). ANI analysis was done using anvi-compute-genome-similarity from anvi’o and can be found within supplemental table 4. ANI relatedness was set with species boundaries of ANI >95% and genus boundaries of >75% (27).

A near full length *16S* rRNA phylogenetic tree was curated using BV-BRC RAST and NCBI databases. Genomes and MAGs which lacked informative *16S* rRNA gene sequences were excluded from analysis. Sequences were aligned using the BV-BRC gene tree service, producing an alignment of 1,560 nucleotide positions. A maximum-likelihood phylogeny was inferred using RaxML v8.2.12 (31) under general time-reversible model with gamma-distributed rate heterogeneity using 1,000 bootstrap replicates. The tree was rooted using *Alcanivorax borkumensis* ABS183 as the outgroup via iTOL (32).

Specific comparison between *T. haligoni* BB40^T^ and *O. antarcticus* wDCs involved recruiting the 158 contigs onto *T. haligoni* BB40^T^’s genome, where 96 contigs were successfully mapped and re-assembled in Geneious v. 2023.1, resulting in 62 scaffolds with a length of 4.50 Mbp. The genome was then put into the BV-BRC server for annotation and aligned back to BB40^T^ for further annotation. Biochemical pathway comparison of *T. haligoni*’s genome to the 13 *Oceanospirillaceae* species was done using BV-BRC server and manual inspection via BLAST and Geneious v. 2023.1 (33–35). Pathway identification was put into three categories based on the presence, partial, or absence identification of essential genes. Identification of complete pathways required all essential genes to be present, while partially complete lacked one of the required genes and absent pathways lacked all the essential genes of the pathway. The NCBI accession number for the genome for BB40^T^ can be found under CP139482.1, and the genome for *O. antarcticus* can be found under JBBKTX000000000.

### Dinitrogen (N_2_) Fixation Rates

N_2_ fixation rate measurements were obtained under hypoxic and oxic conditions for N_2_, NO_3_^-^ (220 *µ*M) and NH_3_ (200 *µ*M) treatments. Hypoxic conditions were consistent to that of temperature range conditions for *T. haligoni*. N_2_ fixation methods and culture enrichment were done as described in (10). Cultures were grown semi-continuously for over 100 days before N_2_ fixation rate measurements were obtained (Fig S1). Acclimated cultures were grown in 250 mL glass culture vials, with modified ASW f/2 + 300 *µ*M phosphate with 6.4 mM carbon (99% purity; Fumarate, Thermo Scientific) and respective N compounds. All bottles were crimped sealed, and hypoxic cultures flushed with helium to purge out oxygen. Helium was used instead of N_2_ so as not to dilute the ^15^N_2_ isotope provided to the culture. Enriched and natural abundance media were added to respective culture bottles for a 5% final enrichment and left to incubate for 24-hours. Harvesting of cultures for POC/PN were collected according to (10). Background ^15^N_2_ (28-, 29- and 30-N_2_) for each replicate was collected in 12 mL triplicate exetainers poisoned with mercury chloride and analyzed on the membrane inlet mass spectrometer (MIMS, Bay Instrument^©^; (36)). Water samples were pumped through a stainless-steel tubing connected to a gas-permeable membrane within the vacuum inlet, both tubing and membrane were immersed in a thermo-regulated bath (6°C) to ensure a constant sampled water temperature at the membrane level. Water vapour was removed from the gas stream by a cryotrap immersed in liquid nitrogen. Oxygen (O_2_) was then removed from the gas stream using a heated copper reduction column (600 °C) to prevent interferences with N_2_ isotopic species and ensure stable ion source conditions. A second cryotrap was used to remove eventual CO_2_, CO and water vapor generated in the reduction column before entering the Pfeiffer Vacuum quadrupole mass spectrometer. Calibration was achieved by sampling a 1 L spherical flask containing *ca.* 500 mL of filtered *in situ* seawater immersed in the same thermoregulated bath containing the membrane inlet. Calibration factors for standard sample were computed based on solubility equations from (37). N_2_ fixation rates were calculated using equation 2:

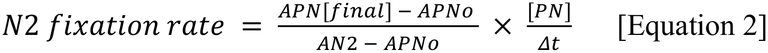

Where *A* is the atom percent of the particulate N (PN; *μ*mol L^-1^) for enriched (*A*PN_final_), natural abundance (*A*PN_o_) cultures and the dissolved background enriched ^15^N_2_ pool (*A*N_2_; %) of the artificial seawater after the incubation period (Δ t; days). Limits of detection for N_2_ fixation rates (nmol N L^-1^ day^-1^ and nmol N cell^-1^ day^-1^) were calculated according to (38) and can be found in supplemental material 3.

### Data Analysis and Availability

Future studies can find the isolated culture under *T. haligoni* BB40^T^, located at the Bigelow NCMA, UBC culture deposit and upon request. The *16S* rRNA sequence for *T. haligoni* BB40^T^ can be found under accession number PZ735692 via NCBI, while the full genome of the isolate can be found under the submission number CP139482.1. All graphical and statistical analysis were conducted in R v.2.2.1 via R studio (R Core Team; RStudio Team 2020). Heatmaps were generated using heatmaply (39), and growth rate curves fitted using standard ggplot2 packages “geom_smooth” with method = “loess, span = 1” (40). Statistical tests for N_2_ fixation rates were standard two-way ANOVA, using the “aov ()” function in baseline R. Unpaired student’s T-tests were used to compare growth rate differences between N sources at specific temperatures. The significance cutoff value for all statistical tests was a p-value less than 0.05 (Table S1). Global mapping of *nifH* distribution was plotted from the following packages: “geom_map”, “rnaturalearth” (41), “geom_sf”, “ggspatial” (42), and “plotly” (43). Genome comparison between *T. haligoni* BB40^T^ and *O. antarcticus* was done in Proksee (44), with BLAST alignment and FASTANI comparison (45).

## Results and Discussion

### Morphology and Physiology

Comparison of *T. haligoni* BB40^T^ to other Oceanospirillales species showed that *T. haligoni* BB40^T^’s genome size is similar in size (4.37 Mbp) to other ORT species and contains a similar GC content (53.4%; Table 1) to that of close relatives. The preferred pH and pH tolerance range of *T. haligoni* BB40^T^ has yet to be tested, however, successful growth for oxic conditions is achieved at pH 8. Lastly, BB40^T^ demonstrates the ability to grow on tween 20, and 80 substrates, which is typical of most Oceanospirillales species given the capability for lipid and hydrocarbon degradation ((18,19,46); Table 1).

**Table 1.** Physiological comparisons of isolated Oceanospirillales species. Isolates were selected based on relatedness to strain BB40T, the availability of data, representation across literature and isolation of cultures. Note that results which have not been determined are indicated by nd. strain BB40T (Rose et al., 2024), *Oceanobacter antarcticus* (Cheng et al., 2025), *Thalassolituus olievorans* (Yakimov 2004), *Teredinbacter turnerae* (Distel et al., 2002), *Parathalassolituus penaei* (Chen et al., 2023), *Marinobacterium litorale DSM 6294* (Bowditch et al., 1984), and *Oceanobacterium kergeii* (Satomi et al., 2002).

| **Variable** | ***T. haligoni*** | ***O. antarcticus*** | ***T. oleivorans*** | ***T. turnerae*** | ***P. penaei*** | ***M. litorale*** | ***O. kriegii DSM 6294*** |
|---|---|---|---|---|---|---|---|
| **Source** | This study | Cheng et al., 2025 | Yakimov et al., 2004 | Distel et al., 2002 | Chen et al., 2023 | Bowditch et al., 1984 | Satomi et al., 2002 |
| **Isolation Source** | marine | marine | marine/sediment | bivalve gills | marine | marine | marine |
| **GC content (%)** | 53.4 | 53.4 | 53.2 | 49 - 51 | 55.4 | 55 | 55 |
| **Genome size** | 4.37 | 4.58 | 3.92 | 5.3 | 2.2 | 3.4 | 4.5 |
| **Growth rate (per day):** |  |  |  |  |  |  |  |
| **NO<sub>3</sub><sup>-</sup> oxic** | 5.8 ** | nd | nd | nd | nd | nd | nd |
| **NH<sub>3</sub> oxic** | 5.2 ** | nd | nd | nd | nd | nd | nd |
| **N<sub>2</sub> hyoxic** | 0.38** | nd | nd | nd | nd | nd | nd |
| **NO<sub>3</sub><sup>-</sup> hypoxic** | 3.8** | nd | nd | nd | nd | nd | nd |
| **NH<sub>3</sub> hypoxic** | 3.8** | nd | nd | nd | nd | nd | nd |
| **Cell shape** | rod | rod | rod and coccoid bodies | rod | rod | rod | rod |
| **Diameter (um)** | 0.5 | 0.5– 0.8 | 0.32 - 0.77 | 0.4 - 0.6 | 0.7 - 1.1 | 0.65 | nd |
| **Length (um)** | 1.5 - 6 | 1.3 – 2.1 | 1.2 - 3.1 | 3 to 6 | 2.2 - 3.4 | 1.6 | nd |
| **Flagellation** | polar (1-2) | Polar (2) | 1 to 4 | 1 | 2 | 1 | 1 |
| **PHB inclusions** | + | nd | - | nd | + | + | + |
| **AlkB gene** | + | - | - | nd | - | + | + |
| **Catalase** | nd | + | + | nd | + | nd | + |
| **Temperature range (T opt) (deg C)** | 4-40 (25)<sup>a</sup><br>4-40 (27-30)<sup>b</sup> | 4 – 40 (10 -15) | 4 - 30 (20 -25) | 30 – 35* | 4 - 30 (25 - 28) | 8 – 42 (20) | 25* |
| **NaCl range (w/v)** | nd | 0 – 4 % (0.5 – 1) | 0.5 - 5.7 % (2.7) | 0.3 | 0 - 4 % (2%) | 1 – 7.5 % | nd |
| **PH range** | 8* | 4- 8 (7) | 7.5 - 9 (8) | 8.5* | 6 - 9 (7) | 5 – 12 (9) | nd |
| **DNRA** | - | - | - | nd | + | + | + |
| ***nifH*** | + | + | - | + | + | - | - |
| **Growth substrate:** |  |  |  |  |  |  |  |
| **Tween 80** | + | nd | + | nd | + | nd | nd |
| **Tween 20** | + | nd | + | nd | nd | nd | nd |
| **Acetate** | + | nd | + | nd | nd | + | nd |
| **Glucose** | - | - | - | nd | nd | + | - |
| **Fumarate** | + | nd | - | nd | nd | nd | nd |
| **Malate** | + | + | - | nd | nd | + | + |

*a: c*ardinal temperatures of N<sub>2</sub> culture
*b*: cardinal temperatures of fixed N cultures
\*: indicates the only condition(s) tested
\*\*: indications T<sub>max</sub> growth rate

TEM and SEM images of exponentially growing *T. haligoni* BB40^T^ cells revealed minimal physiological difference between fixed N and N_2_ conditions (Fig 1). SEM pictures showed that all cells displayed a wrinkle cell surface and possessed flagella. SEM pictures revealed circular bodies throughout the cell. Size ranges across the treatments ranged between ∼1.5 – 6.0 um in length, with N_2_ conditions having longer rod-like cells (Fig 1A). All growth conditions led to the secretion of an extracellular polymer, with N_2_ conditions secreting a thicker film-like substance (Fig 1A-C; Fig S3). TEM images and gas chromatography also demonstrate the isolate has the ability for polyhydroxybutyrate (PHB) synthesis (Fig 1, Fig S4). While PHB accumulation differences were not seen between the N sources (oxic and hypoxic), there was a difference in PHB accumulation when the isolate was grown on different carbon sources in the presence of NO_3_^-^ (Fig S5). Growth on the mixed carbon cocktail (glucose, fumarate, succinate, lactate and acetate) in comparison to a singular carbon source (fumarate) had the C storage molecule occupy most of the cellular space (Fig S5).

**Fig 1.**
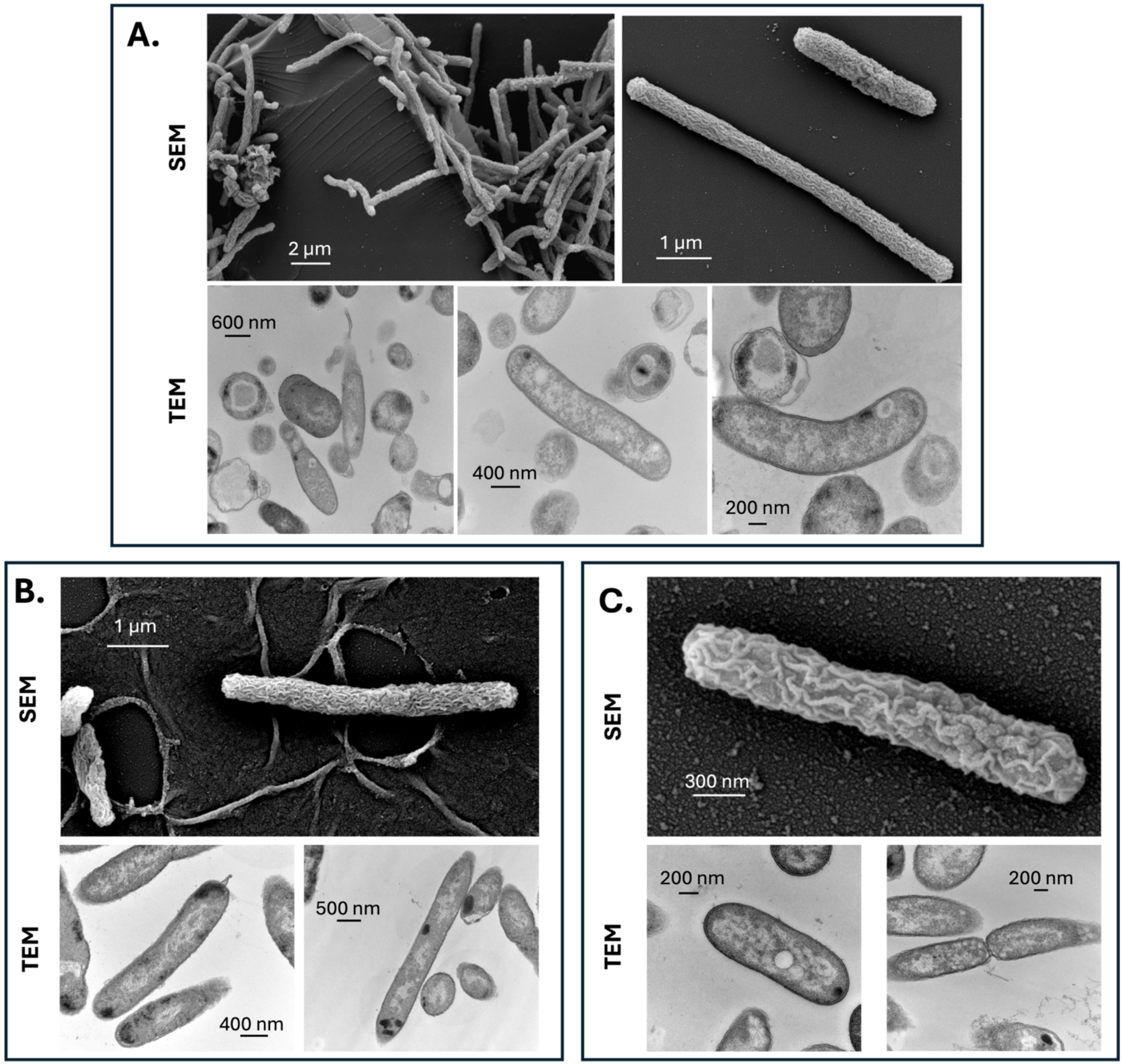
*Thalassolituus haligoni* sp. nov. BB40T scanning (SEM) and transmission (TEM) electron microscopy at 15 °C under N2 **(A),** NH3 **(B)** and NO_3_^-^ **(C)**. Images collected under different magnification and electron voltage. Cells were collected in an exponential phase. Details for electron microscopy magnifications can be found in Data S2. SEM microscopy details are as follows: N2 (EHT= 5 kV, signal A= SE2, WD = 11.4 mm, mag = 14.79 KX); NH3 (EHT= 5 kV, signal A= SE2, WD= 11.4 mm, mag – 13.9 KX); NO_3_^-^ (EHT= 5 kV, signal A= SE2, WD= 11.4 mm, mag = 26.76 KX). TEM microscopy details are as follows: N_2_ (*R*: HV= 80 kV, mag= 40 KX; *M:* HV= 80 kV, mag= 40 KX; *L:* HV= 80 kV, mag= 50 KX); NH_3_ (L: HV= 80 kV, mag= 40 KX *;R:* HV= 80 kV, mag= 30 KX); NO_3_^-^ *(L*: HV= 80 kV, mag= 60 KX;*R:* HV= 80 kV, mag= 50 KX); with abbreviations *M=*middle photo, *R=* right photo and *L*= left photo.

Thermal performance curves for BB40^T^ across the five N conditions demonstrated growth rates ranging between 0.10 (± 0.01) – 5.80 (± 1.19) per day ^-1^ and were significantly higher in fixed N than N_2_ treatments, with oxic conditions having the highest growth rate overall (Table 1; Fig 2; Fig S2). The size, shape and temperature range of the isolate is also characteristic of *Oceanospirillales* species, with bi-polar flagella, rod shape appearance, and growth between 4 - 40 °C (Table 1; Fig 2; (18)). Hypoxic growth conditions maintained a flatter thermal performance between 4 - 35 °C but were still able to maintain growth at 40 °C (Fig 2). Reasoning for maintained growth at 40 °C in hypoxic fixed N cultures remains unknown, however, given detection of the isolate within oxygen minimum zones (OMZs) at deeper waters, it could be a physiological adaptation for growth near hydrothermal vents (10,47–49). In contrast, oxic conditions demonstrate a more traditional growth performance curve, having faster growth rates with increasing temperature, up to a maximum (i.e., 30 – 35 °C), and after which, collapse at 40 °C (Fig 2). The difference in higher temperature optima between hypoxic and oxic conditions is predicted to be the result of different growth limitation bottlenecks (i.e., reactive oxygen species in oxic cultures cannot be managed at higher temperatures vs. energy supply is lower in hypoxic cultures (50)). When N demands of the cell are not met however, and BB40^T^ is required to fix N_2_ (i.e., N_2_ hypoxia), the thermal performance shifts to a temperature growth maximum between 21 - 25 °C (Fig 2). We predict that this significant shift in thermal performance is reflective of a species-specific thermal preference for nitrogenase such as seen in *Klebsiella pneumoniae* (Fig 3.2; (51,52)). Given the cells are restricted by fixed N during true diazotrophic conditions, the main bottlenecks for the N_2_ hypoxic conditions appears to be nitrogenase synthesis, and N_2_ fixation upkeep.

**Fig 2.**
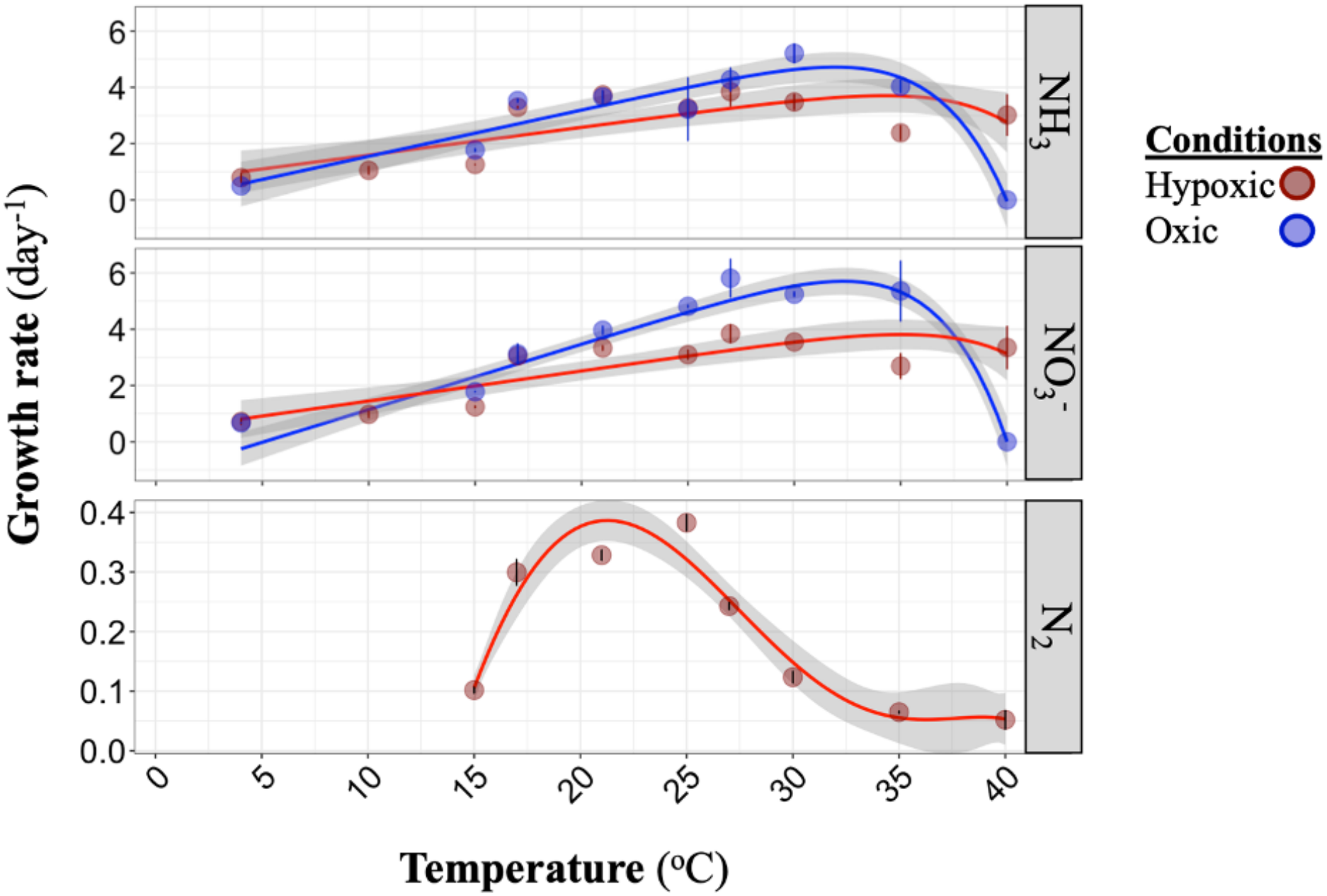
Average growth rates of strain BB40T ± 1 standard error under various fixed N sources (220 µM NO_3_^-^, 200 µM NH_3_, and N_2_) and temperatures (°C). Error bars indicate the average of biological triplicates. Growth curves for 15 ° C data can be found in supplemental. Growth rate calculations can be found in supplemental material 3 and growth curves can be found in supplemental Fig S2.

**Fig 3.**
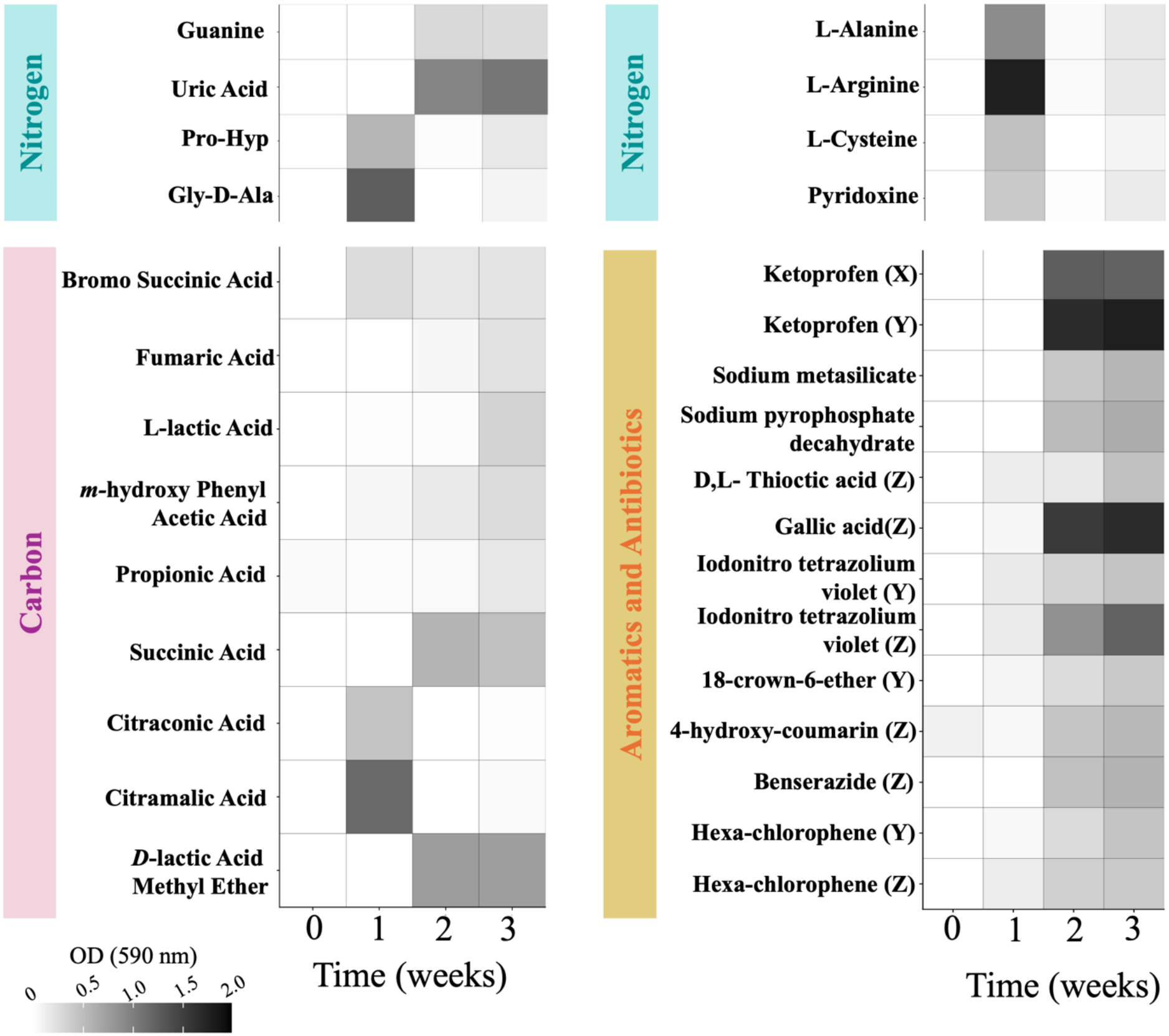
Cell growth over time (weeks) of BIOLOG PM plates of positive conditions of strain BB40T. Cells measured under optical density (OD) 590 nm. C plates (pink) were supplemented with 220 µmol NO_3_^-^, with the described sole C source from the plate. N plates (green) were supplemented with 6.4 mM C (fumarate), with the N source provided by the plate well. Aromatic and Antibiotic plates only included the inoculating ASW media (Data S2) while each well within the plate was the sole media used for growth.

**Fig 4.**
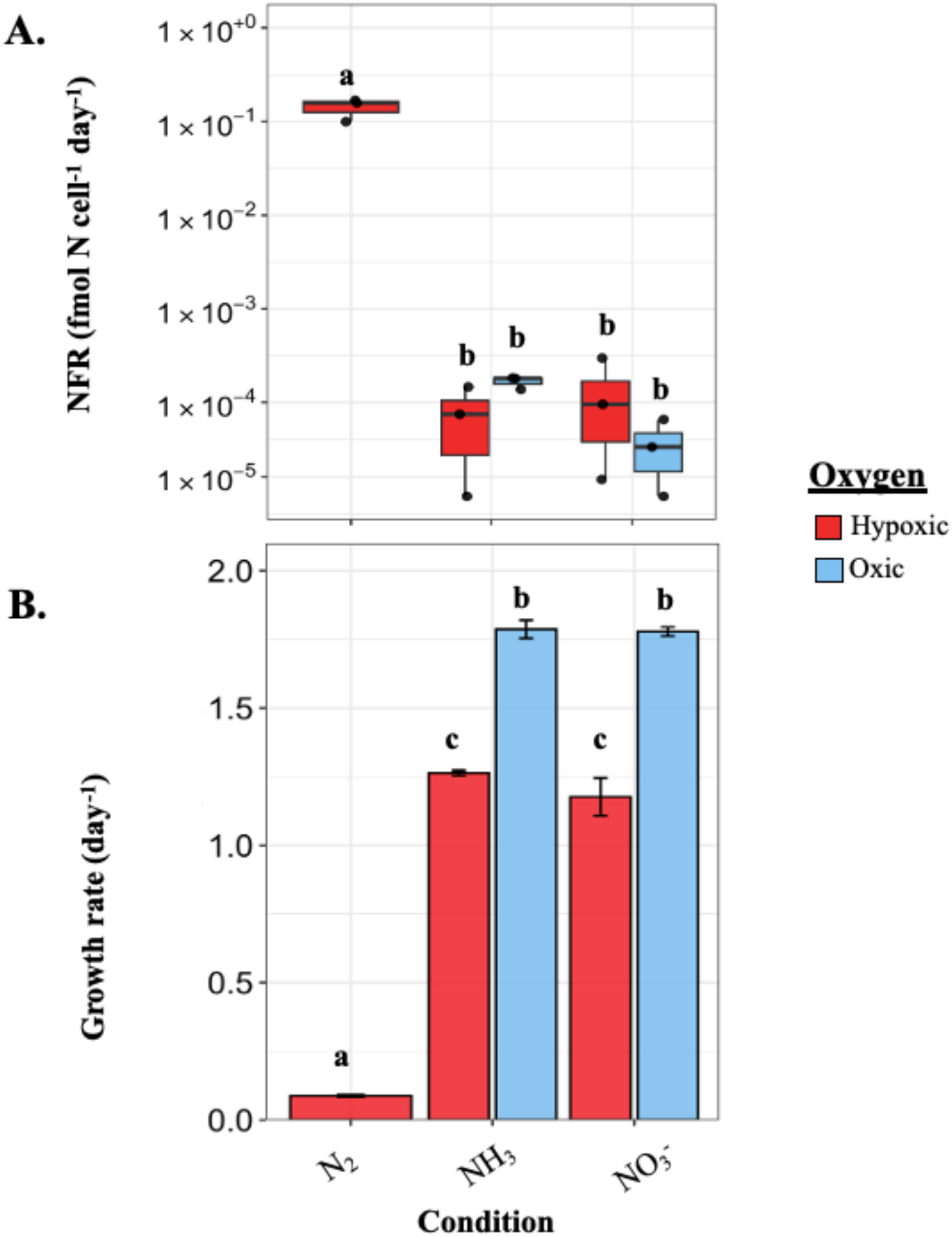
N_2_ fixation and growth rates of five conditions for strain BB40T at 15 °C. **A)** Average N_2_ fixation rate measurements (fmol N cell^-1^ day^-1^) ± 1 standard error of strain BB40T under various O_2_ (hypoxic, oxic) and N conditions (N_2_, NO_3_^-^, NH_3_). **B)** Growth rates (per day) of strain BB40T under hypoxic and oxic conditions for N_2_, NO_3_^-^ and NH_3_ sources. The statistical test conducted was the standard student’s T-test, with p-value = 0.05. Note the y-axis is displayed on log scale for panel A. Any shared letters were determined to not be statistically different.

**Fig 5.**
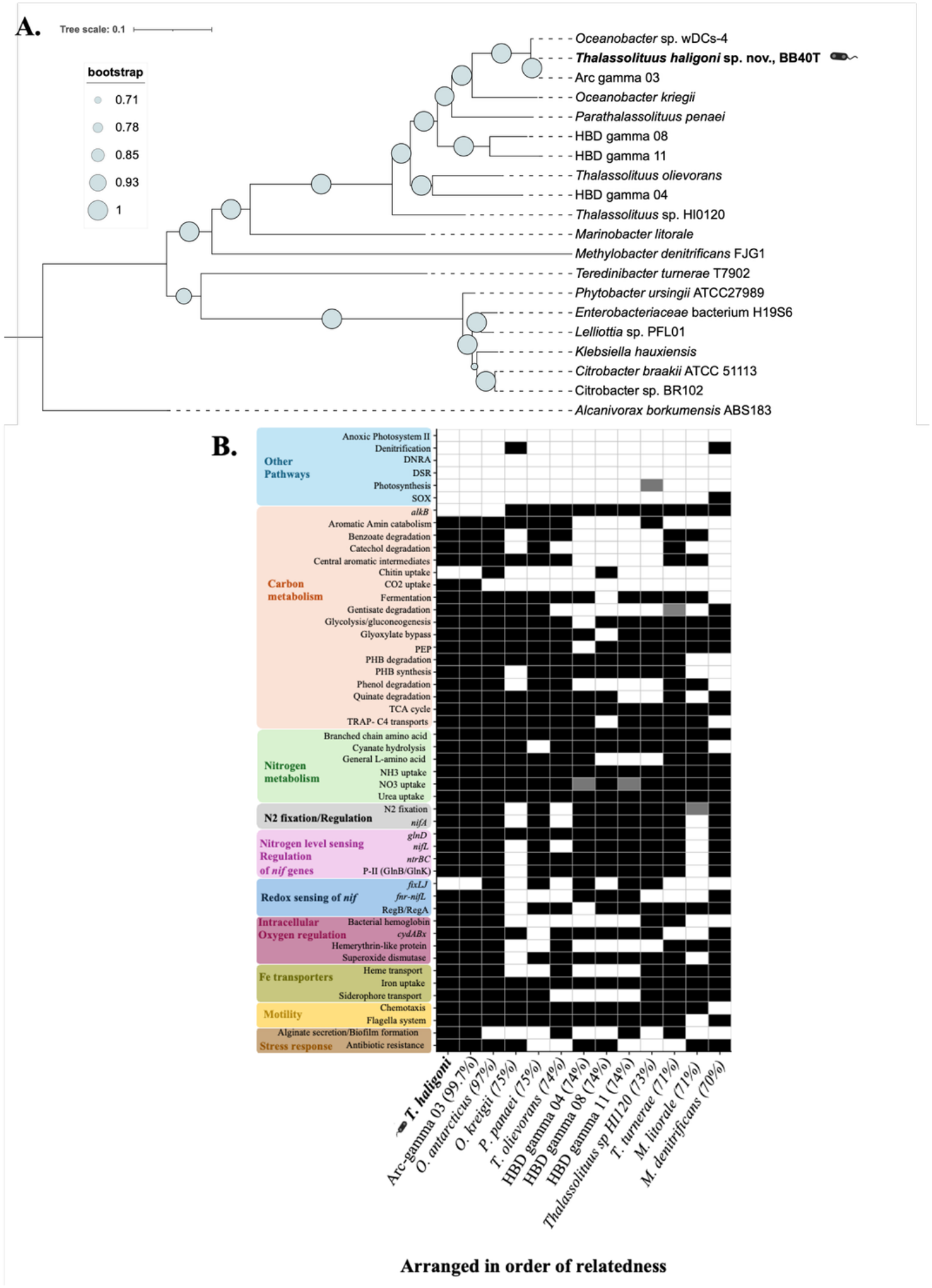
Genome and phylogenomic comparison of Oceanospirillales. **A)** Phylogenomic tree of 13 closely related genomes. **B)** Genome comparison of closely related species to strain BB40T. Genomes are ordered based on relatedness to strain BB40T and have the ANI relatedness indicated in brackets. Pathways of interest were selected based on evidence provided from Turk-Kubo et al., (2022), strain BB40T laboratory growth and common N_2_ fixation requirements (i.e., O_2_ levels and Fe). Complete pathways displayed in black, partial pathways displayed in grey and absent pathways displayed in white. Pathway confirmation was determined through the RAST server, KEGG, Geneious, BLAST and evidence already provided from Turk-Kubo et al., 2020, Fig5.

Substrate verification on PM Biolog assay plates showed a wide range of phenotypic capabilities for the isolate. C plates (PM 1 and 2a) had positive reduction of tetrazolium dye on C_4_ dicarboxylic acid substrates, Lactic acid, Phenyl Acetic acid, Propionic acid, and D-lactic acid methyl ester. N plates (PM3) were positive successful for Uric acid, and Guanine, and nutrient supplements, while PM5 plates were positive for L-Alanine, L-Arginine, L-Cysteine, and Pyridoxine (Fig 3.3). Peptide N sources (PM 6-8), had two positive wells for Pro-Hyp and Gly-D-Ala. Other plates testing for chemical sensitivity (PM 14-20) such as antibiotic resistance, heavy metals and phenolic compounds, were positive for Hexa-chlorophene (Y and Z), Ketoprofen (X,Y), Gallic acid (Z), Benserazide (Z), 4-Hydroxy-coumarin (Z) and 18-crown-6-ether (Y) (Fig 3.3). Substrate preference was supplemented with standard laboratory culture growth tests which further expanded the diversity of BB40^T^ (10).

N_2_ fixation rates (NFR) for *T. haligoni* BB40^T^ were confirmed in (10), and expanded upon in this study (Fig 3.4A). NFR measurements in this study confirmed the isolate can fix N_2_ under a range of N and O_2_ conditions (9.34 × 10^-6^ to 1.4 × 10^-1^ fmol N cell^-1^ day^-1^; Fig 3.4A) at the expense of reduced growth rate (Fig 3.4B). Hypoxic N_2_ conditions were over 3 orders of magnitude higher than fixed N conditions (hypoxic or oxic), demonstrating levels between Gamma-A and *S. castanea* (53,54). To date, the NFRs measured for *T. haligoni* are the highest reported for any free-living NCD, suggesting that *T. haligoni* BB40^T^ is a globally significant contributor to marine N_2_ fixation. In addition, the significant reduction in growth rate (90%) for N_2_ hypoxic conditions (Fig 3.4B) at the expense of N_2_ fixation coincides with other diazotrophic studies, due to the energy demands of the process and need to meet N-demands of the cell ((55–57). Within the fixed N treatments, none of the NFR rates were significantly different from each other, regardless of fixed N or O_2_ source (Fig 3.4A; supplemental material 3). While this result is counterintuitive with respect to energy uptake between NH_3_ vs. NO_3_^-^, it suggests either an alternative use for nitrogenase or maintenance of nitrogenase at a basal level to elicit a rapid response to external conditions. The NFR in NH_3_ conditions for example, could result from slower NH_3_ uptake as a mechanism to manage toxicity, where reduced uptake of fixed N is compensated for by increased N_2_ fixation in low O_2_ conditions (58).

### Genomic characteristics

The complete genome (53.42 mol%) of *T. haligoni* BB40^T^ was obtained and characterized via RAST with 462 subsystems, 4,019 CDS genes and 61 RNA genes (9 rRNA and 52 tRNA) (10). The genome was determined to be 99.77% complete, with a contamination of 1.07 (supplemental material 4). The genome contains 132 CDS for cell wall and capsule formation, 108 CDS for motility and chemotaxis, 171 CDS for Fatty acids, lipids and isoprenoids, 69 CDS for nitrogen metabolism, 127 CDS for metabolism of aromatic compounds, 216 CDS for carbohydrate metabolism, and 154 CDS genes for oxygen metabolism (*see supplemental material* (10)).

Cell wall genes display all genes required for gram negative cell wall formation and contain genes for alginate secretion (capsular and extracellular polysaccharides). Motility genes represent a complete prokaryotic flagellar system (including *flhABF, fliIRMN, flgDH, motAB and fleNF*), with evidence of genomic redundancy. The isolate also possesses chemotaxis genes (*cheAVY*), indicating the capability for directional movement. Nitrogen metabolism indicates a full suite of *nif* genes (26 CDS), containing *nifHDKENTBQOXUSVWZMAL* and accessory genes *nifX-2, FrdN, sufA, nafY, LRV, CysE, IscA.* Surrounding the *nifHDK* cluster, *Avin2460,* and various hypothetical proteins are found, indicating possible involvement in N_2_ fixation, however, further investigation is required. N metabolism within the genome also shows NH_3_ assimilation, cyanate hydrolysis and NO_2_ / NO_3_^-^ ammonification. Carbohydrate metabolism contains several subcategories, including CO_2_ fixation (3 CDS; RuBisCO transcriptional regulator, and 2 Carbonic anhydrase), fermentation (58 CDS; Acetyl-CoA to Butyrate, Lactic acid fermentation, Butanol biosynthesis and Acetolactate synthase subunits), organic acid degradation, and central carbohydrate metabolism. Membrane transport within the genome contains the ability for TRAP C_4_ uptake, confirming the ability for growth on C_4_ dicarboxylic acids, and also includes glyoxylate uptake, heme transports, Nickel, Magnesium, Cobalt, and Copper transporters, and phosphate transport.

Vitamin production pathways present within the genome include thiamin biosynthesis, biotin (*bioMNY; bpr; bioC1BxBAFGD*), folate biosynthesis (*folEQXKP; dhfSR; dhnA; pabA-bc;* dTMP and several folate transporters), molybdenum cofactor biosynthesis (*moaABCED; mobAB),* and coenzyme B12 biosynthesis (*cbiJTCAP, cobDUTCS, pduXS*). Vitamin transports within the genome include Mo permease and ATP-binding protein, vitamin B12 ABC transporter, TonB membrane transports (*BtuB*), Fe^3+^ and Fe^2+^ transports, hemin transports, and ferric siderophore transport. With the genomic capability for B12 biosynthesis and membrane uptake, the examination of *T. haligoni* BB40^T^’s role as a B12 producer or scavenger has yet to be investigated, however, could suggest an interesting link between NCDs and cobalamin production (25,59–61).

O_2_ metabolism of BB40^T^ includes oxidative stress mechanisms – protection from ROS, glutathione; redox and non-redox reaction – dioxygenases, and anaerobic respiratory reductases in electron accepting reactions [Flavodoxin, Ferredoxin, Coenzyme F420, Vanillate O-demethylase oxidoreductase, ubiquinone oxidoreductase, Fe-S-cluster containing hydrogenase, and Arsenate reductase]. Stress mechanisms within the isolate are also variable, containing bacterial hemoglobins, flavohemoglobin, C starvation mechanisms, osmotic stress and detoxification transporters, heat shock (*dnaK* gene cluster), and cold shock (*cspA* family) pathways.

The genome also contains remnants of integrons (12 CDS with redundancy) and glutaredoxin phage associated genes. In particular, the presence of integrons suggests possible antibiotic resistance acquired via phage DNA while glutaredoxin (*grxR*) being linked to helping with oxidative stress and nitrogen management within the cell (62–64). Although phage–NCD interactions remain poorly understood, the presence of integrons and phage-associated genes may be key to the ecology of *T. haligoni* in dynamic environments (i.e., OMZs and particle-associated habitats), though their functional roles remain unexplored. Future industrial applications for the isolate are also found within the genome, with the identification of aromatic degradation for oil intermediates (quinates, salicylates, phenols and benzoates), alkane oxidation of medium chain alkane groups (C_5_ – C_22_) (65–67), and possible plastic alternatives through the optimization of PHB metabolism (68,69). While yet to be tested, BB40^T^ could also play a significant role in deep sea C and N cycling given the presence of alkane oxidation genes, temperature tolerance in hypoxic conditions and detection of *nifH* ASVs in OMZs and hydrothermal vents.

### Phylogenomic Analysis and Genome Comparison

Phylogenomic relationships to *T. haligoni* BB40^T^ demonstrate it belongs within an *Oceanospirillales* clade of uncultured diazotrophic MAGs (i.e., Arc-gamma-03), collected from the TARA expedition ((10,12,13). This placement was broadly supported by the 16S rRNA gene phylogeny (fig S6). In addition, *T. haligoni* BB40^T^ shares a close relationship with *Oceanobacter antarcticus* wDCs (ANI 97.16 %)*, Parathalassolituus penaei* (ANI 74.80 %), and *non*-diazotrophic species: *Oceanobacter kriegii* (ANI 75.05%), and *Thalassolituus* oleivorans (ANI 74.02%; Fig 3A; supplemental material 4). The closest relative, Arc-gamma-03, although uncultured, has 99% genomic identity with *T. haligoni* BB40^T^, and contains 100% identical *16S* rRNA and *nifH* sequences, suggesting a closely related arctic ecotype ((10); supplemental material X), whereas, the second closest relative, *O. antarcticus* shares 99.93% identity to *16S* rRNA and 99% *nifH* identity (supplemental material 4). Given the limited number of diazotrophic Oceanospirillales isolates, understanding the N_2_ fixation potential and physiology of *T. haligoni* BB40^T^ and other close NCD relatives (i.e., *O. antarcticus* wDCs) enhances our understanding of the ecological function and biogeochemical significance of this lineage within marine C and N cycles.

The degradation of phenolic compounds within this *Oceanospirillales* group shows a patchy presence/absence within the genomes, suggesting an environmental response for the acquisition of these genes (70,71). Characteristic of *Oceanospirillales*, the majority of the genomes display the *alkB* gene, flagella, chemotaxis systems, fixed N uptake and the PHB degradation/synthesis cycle (Fig 3B), however, *T. haligoni*, *O. antarcticus* and ARC-Gamma-03 lack this gene. Examination of *nif* regulation genes appears to be fairly conserved across the taxa, however, more distantly related *M. litorale,* and *O. kiregii* lack the *ntrBC*, *nifL* and PII genes typically seen within *nif* containing genomes. Lastly, most of the genomes do not display the ability for “other pathways”, exceptions to this are seen in *O. kiregii* and *M. denitrificans* (denitrification) (Fig 3.3B).

*O. antarcticus* wDCs (153 contigs; 4.58 Mbp) and *T. haligoni* BB40^T^ (4.2 Mbp) genomes shared 25 RAST categories (81 subcategories; 1232 CDS), with *T. haligoni*, having 25 distinct categories (89 subcategories; 524 CDS) (Data S4), and *O. antarcticus* having 20 distinct categories (23 subcategories; 65 CDS). Of the shared categories with specific annotation (i.e., excluding hypothetical proteins), shared gene similarity was highest within amino acids and derivatives, RNA metabolism, cell wall and capsule formation, cofactors and vitamins, stress response, respiration and N metabolism (supplemental material 4). Specifically, within N metabolism, both genomes contain a full suite of *nif* genes for N_2_ fixation, including *nifHDK* (supplemental material 4). Genes specific to *O. antarcticus* (65 CDS), were mostly involved in conjugative transfer (i.e., nucleoprotein and secretion systems; *trbBCDEFGHIJL)*. Another similarity between the two strains is the lack of nickel (Ni)-Fe hydrogenase associated with N_2_ fixation metabolism. While it remains unclear how diazotrophs which lack the Ni-Fe specific hydrogenase deal with the accumulation of H_2_, suggested hypotheses include: alternate hydrogenases (i.e., bidirectional or uptake hydrogenases), acetogenesis or fumarate reduction (72,73). Given the isolate’s preference for fumarate, this suggests that any increase in H_2_ gas is managed through fumarate reduction, negating the need for a specific N_2_ fixation hydrogenase. Notably, the genome of *O. antarcticus* lacks a bacterial secretion system, a ribose transport system, betaine biosynthesis, while BB40^T^ contains the genomic capacity for these metabolisms.

Physiological comparisons between *T. haligoni* BB40^T^ and *O. antarcticus* wDCs also highlight interesting differences (Table 1; (74)). *T. haligoni* shows growth best between 20 - 30 °C, while *O. antarcticus* is more suited for temperate temperatures (10 - 15 °C). The growth range between the two isolates is similar, however, *O. antarcticus* shows capability of growth up to 40 °C, while *T. haligoni* fails to grow at this temperature in fixed N oxic conditions. More surprisingly, the pH range of *O. antarcticus* was determined to have capable growth from 4 - 8, with an optimum of 7. While the full pH range of BB40^T^ has yet to be determined, preliminary work and successful growth of the isolate estimate a much narrower pH tolerance range, as growth is only successful (to date) between 7.8 - 8.3 in an oxic environment. Examining the distribution between the two species, *O. antarcticus* occupies mostly temperate and polar regions, while *T. haligoni* occupies a more global range (Fig S7;(10)). Given the close genomic relationships between *O. antarcticus* and *T. haligoni* and the global distribution of BB40^T^ these findings highlight the physiological diversity among diazotrophic ORTs and establish *T. haligoni* BB40^T^ as a valuable model for understanding NCD N_2_ fixation and its response to climate change.

## Conclusion

*Thalassolituus haligoni* BB40^T^ was determined to be a facultative anaerobic gram-negative flagellated heterotrophic bacterial diazotroph. The globally distributed isolate shares close relatedness to the polar species, Arc-gamma-03 (99 % ANI) and the recently cultivated *O. antarcticus* (97 % ANI), forming a significant but understudied clade of ORT NCDs (Delmont et al., 2018, Zhang et al., 2025; Fig 3A, Data S5). Laboratory studies of *T. haligoni* BB40^T^ confirm possible growth on a variety of C, N and antibiotic compounds, with a preference for C_4_ dicarboxylic acids. NFR for the isolate suggests a basal background nitrogenase expression, while active N_2_ fixing cultures demonstrate the highest NFRs for a free-living cultured NCD, and could be on a similar magnitude to Gamma-A once single-cell rates are achieved (53). More importantly, successful N_2_ fixation is more impacted by the presence of O_2_, rather than fixed N sources, and is highlighted by the variety of O_2_ protective mechanisms found within the genome (i.e., redox, *cydABx*, alginate secretion) and the inability for N_2_ growth in oxic conditions. Collectively, the isolate demonstrates a wide versatility in its genome and physiology which can be used for anthropogenic cleanup (i.e., bioremediation) and understanding links within biogeochemical cycles. The work provided herein provides an in-depth species description for a globally distributed NCD.

## Supporting information

supplemental material 1 Thalassolituus haligoni v.s. Oceanobacter antarcticus ASV presence/ absence global distribution map.

supplemental material 2 Artificial Seawater Media Recipe

supplemental material 3 Calculated Growth rates and statistical tests for thermal tolerance curves.

supplemental material 4_genome information_ANI

## Acknowledgements

We would like to acknowledge laboratory members Scott Pollara, Megan Roberts, Ella Joy H. Kantor, Nan Chen, and Becca-Stevens Green who have helped with the sampling and instrument setup for this work. We would also like to give a special thanks to Maria Armstrong, and Dr. Christopher Algar who provided resources for the hypoxic growth conditions and Claire Normandeau for training and help with the EA-IRMS.

## Author Contributions

S.A.R., S.L.G.D., E.M.B., and J.LR designed the experiment; J.LR., and E.M.B supervised the experiment; S.A.R., and S.L.G.D., carried out the experiment; S.A.R., and S.L.G.D analyzed the data; S.A.R drafted the original manuscript; S.A.R., S.L.G.D., P.L., G.C., J.T., E.M.B., and J.LR revised the manuscript.

