## supplemental figures for "*Thalassolituus haligoni* BB40^T^, a model species for non-cyanobacterial diazotrophs within Oceanospirillales isolated from a Fjord-like Inlet in Kjipuktuk"

### Supplemental table and Figures

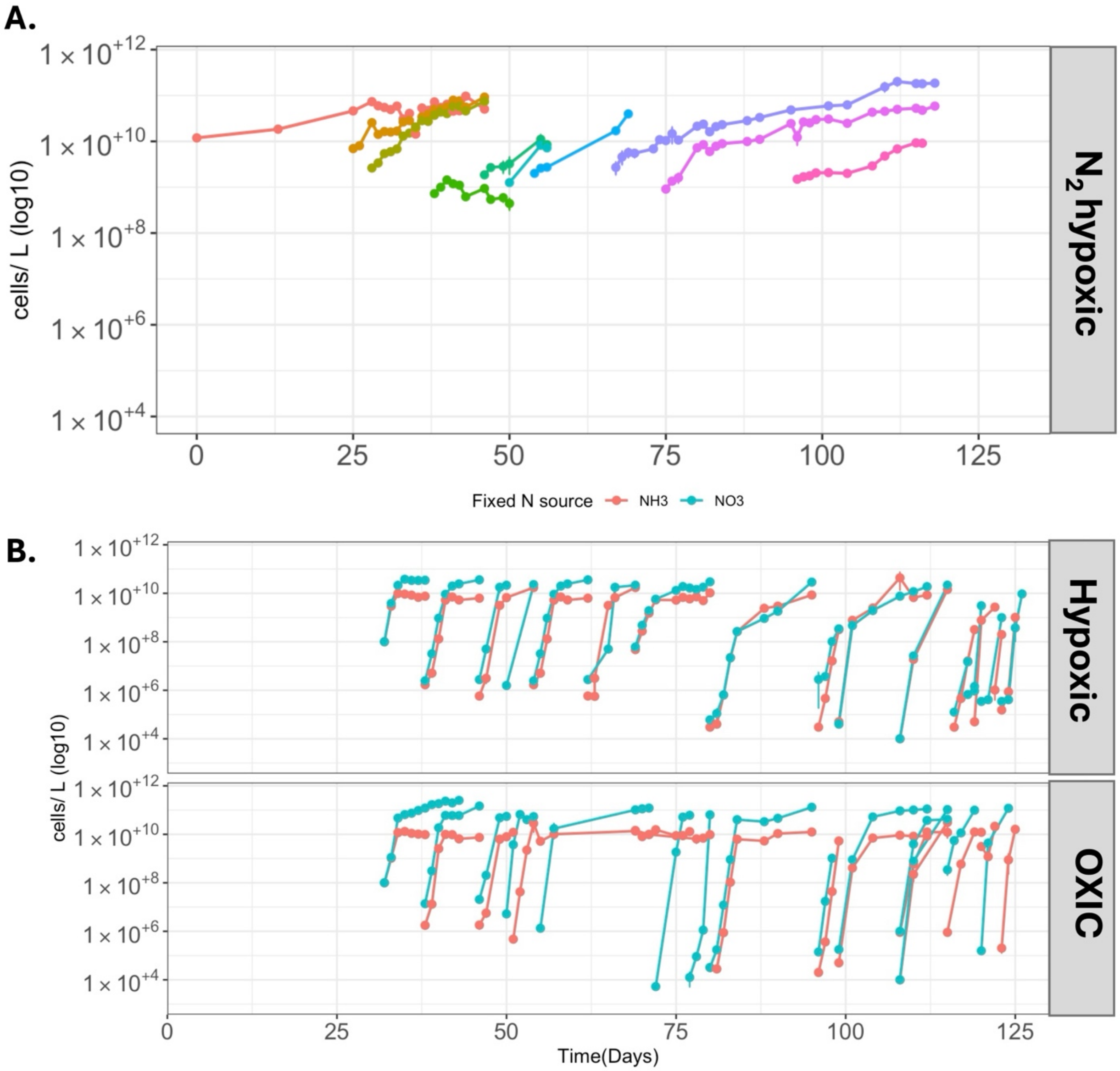


**Fig S1** Average cell density of semi-continuous growth curves of *Thalassolituus haligoni* under $N_2$ at 15 $^o$ C (**A**) and fixed N [$NO_3^-$ (blue), $NH_3$ (Orange); **B**) conditions in hypoxic and oxic treatments. Error bars represent ±1 standard error. Note the different colours in panel A) are $N_2$ transfer cultures.

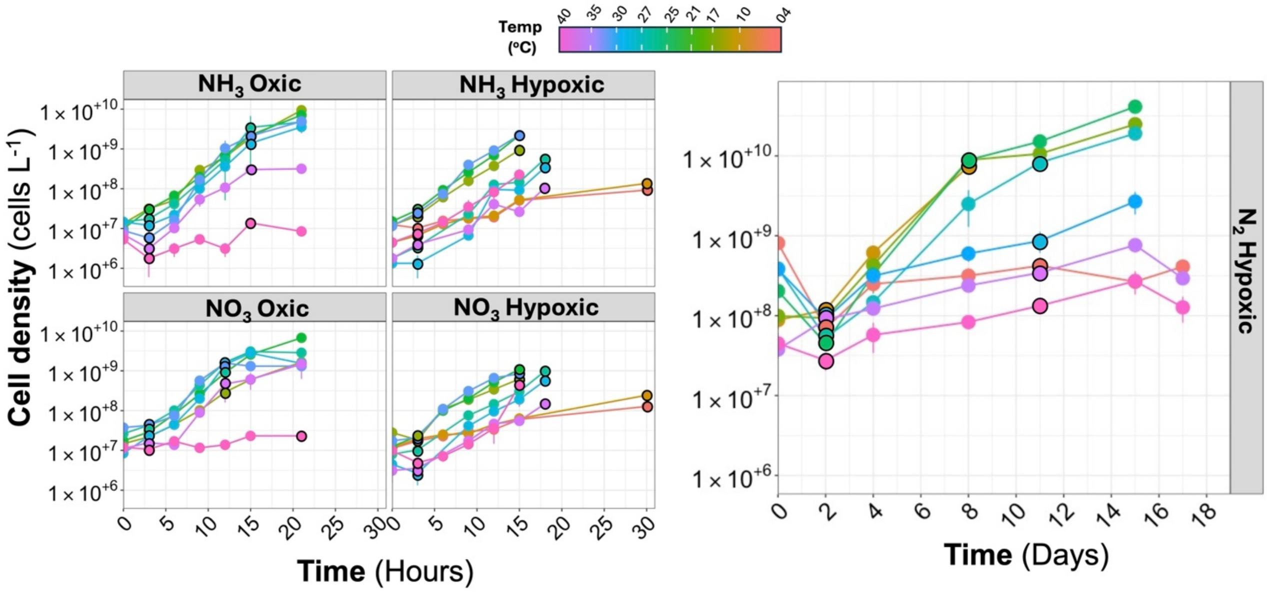


**Fig S2** Growth curves of *T. haligoni* ± 1 standard error under various fixed N sources (220 uM NO3, 200 uM NH3, and N2) and temperatures (deg C). Error bars indicate average of biological triplicates. Black outlined circles indicate T0 and T-final used for calculating growth rate. Growth rate calculations can be found in supplemental material 3.

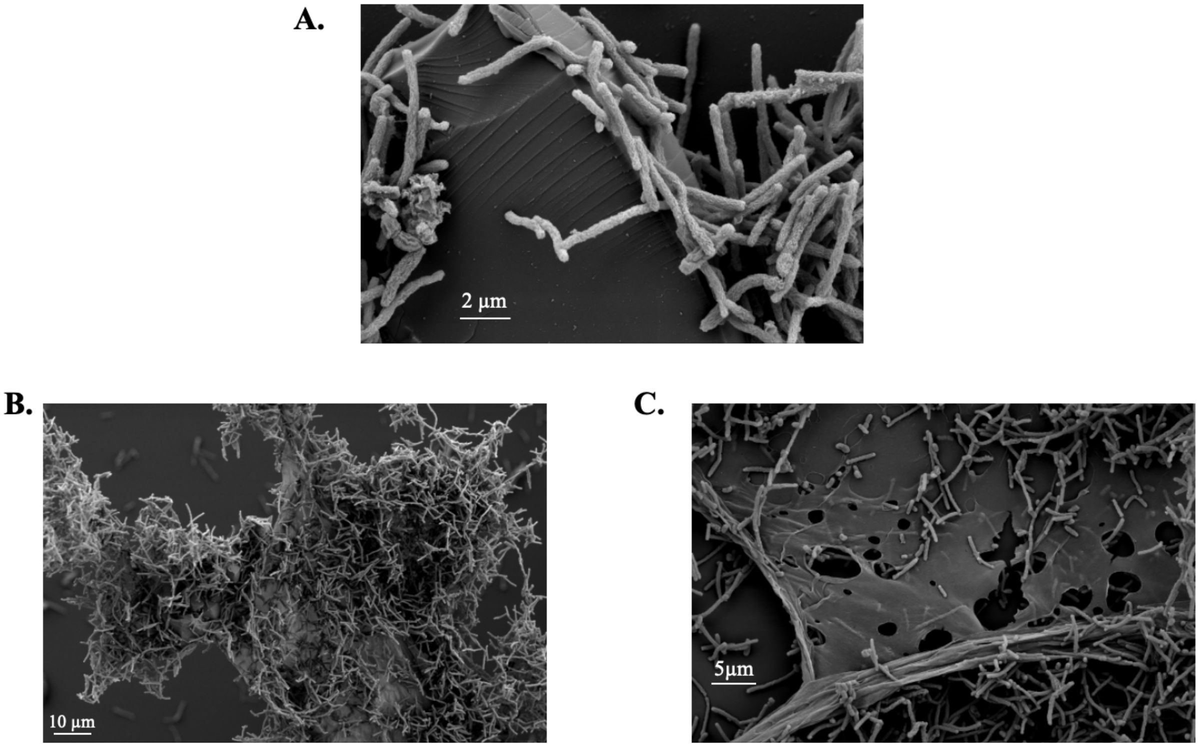


**Fig S3** Scanning electron micrographs of *T. haligoni* under hypoxic $N_2$ (**A**), oxic $NO_3^-$ (**B**) and oxic $NH_3$ (**C**) conditions. Cultures were grown at 17 ºC in modified f/2 ASW with 6.4 mM C (fumarate), under atmospheric $N_2$, 200 µM $NO_3^-$, or 220 µM $NH_3$. Microscopy parameters for images include EHT = 5.00 kV; WD= 11.1 mm; Signal A= SE2. Magnification for A) was set to 5.45 KX; B) 813 X and C) 1.97 KX

S1= PHB S2= PHB coPV S3= sample

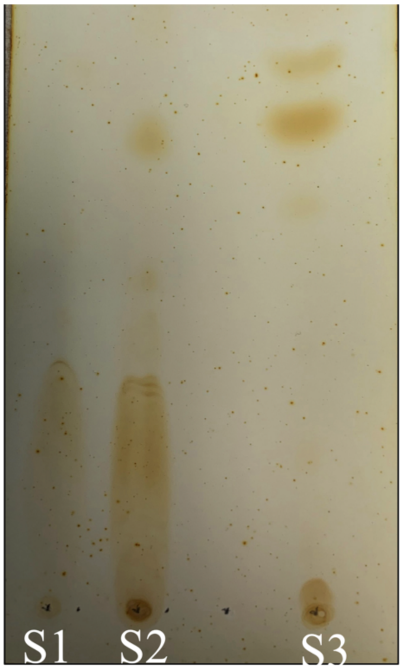


**Fig S4** TLC analysis of PHBs in *T. haligoni* under mixed carbon cocktail and depleted N conditions. TLC conducted by Zhenyu in Tupper building 2020. $R^f$ 0.47, 0.56, 0.65.

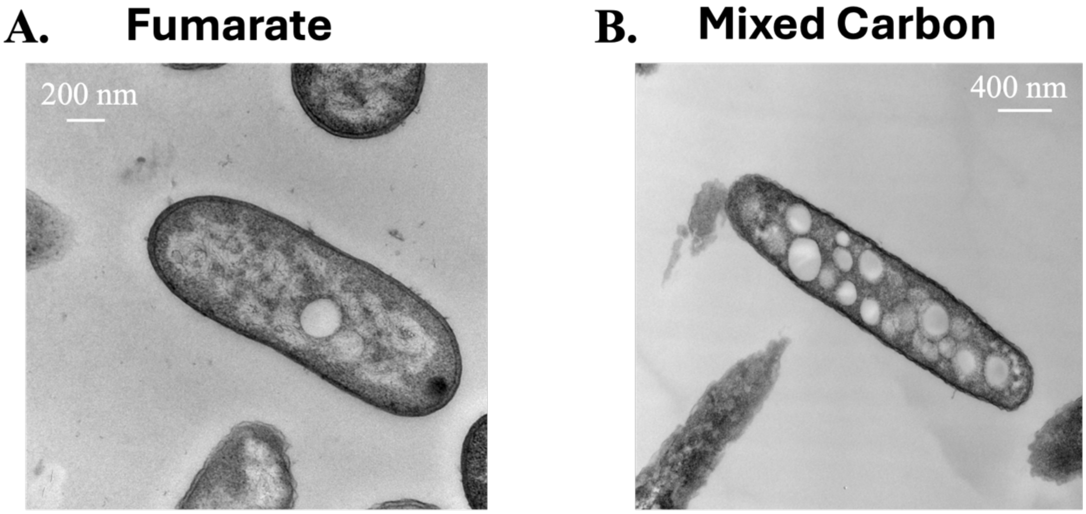


**Fig S5** Transmission electron microscopy of *Thalassolituus haligoni* under different carbon sources (6.4 mM C) [fumarate or mixed carbon cocktail (MCC)], under replete nitrate conditions (220 uM). Cells were grown at 17 ºC in the exponential phase of growth.

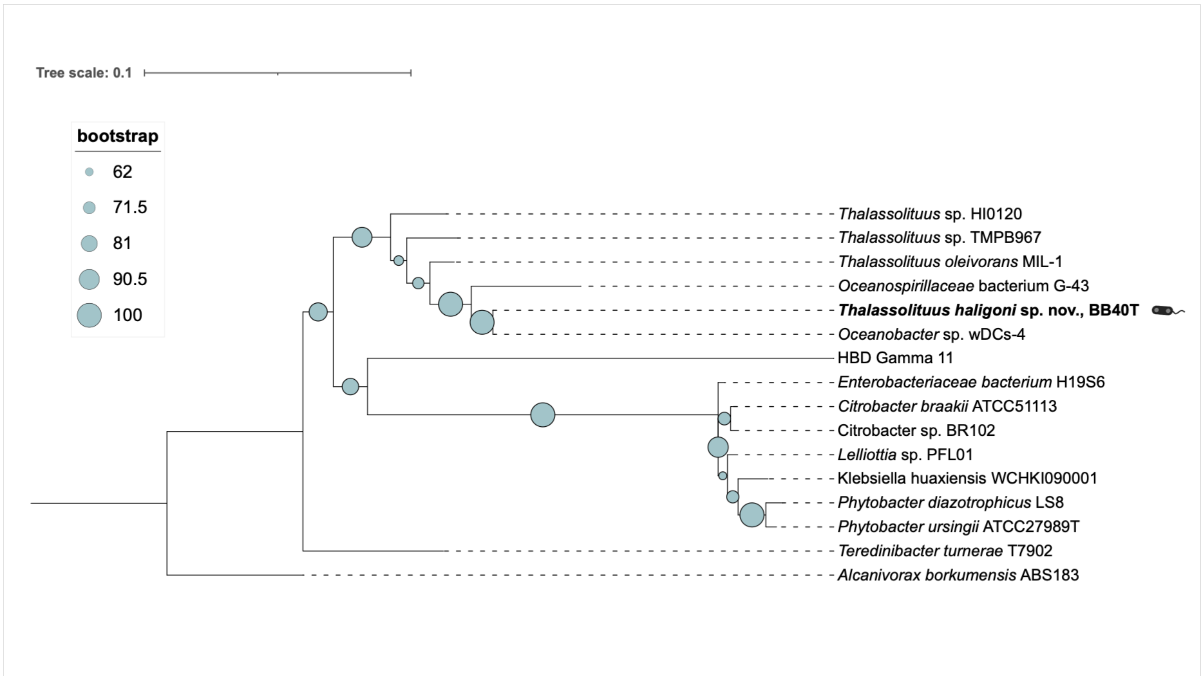


**fig S6 16S rRNA gene phylogeny of *Thalassolituus haligoni* BB40$^T$ and related taxa.** The maximum-likelihood tree was reconstructed from an untrimmed 1,560-position 16S rRNA gene alignment using RAxML v8.2.12 under the GTR+ gamma model. Branch support was assessed using 1,000 bootstrap replicates and is represented by circles proportional to the corresponding support value. The tree was rooted using *Alcanivorax borkumensis* ABS183. The scale bar represents 0.1 nucleotide substitutions per site.

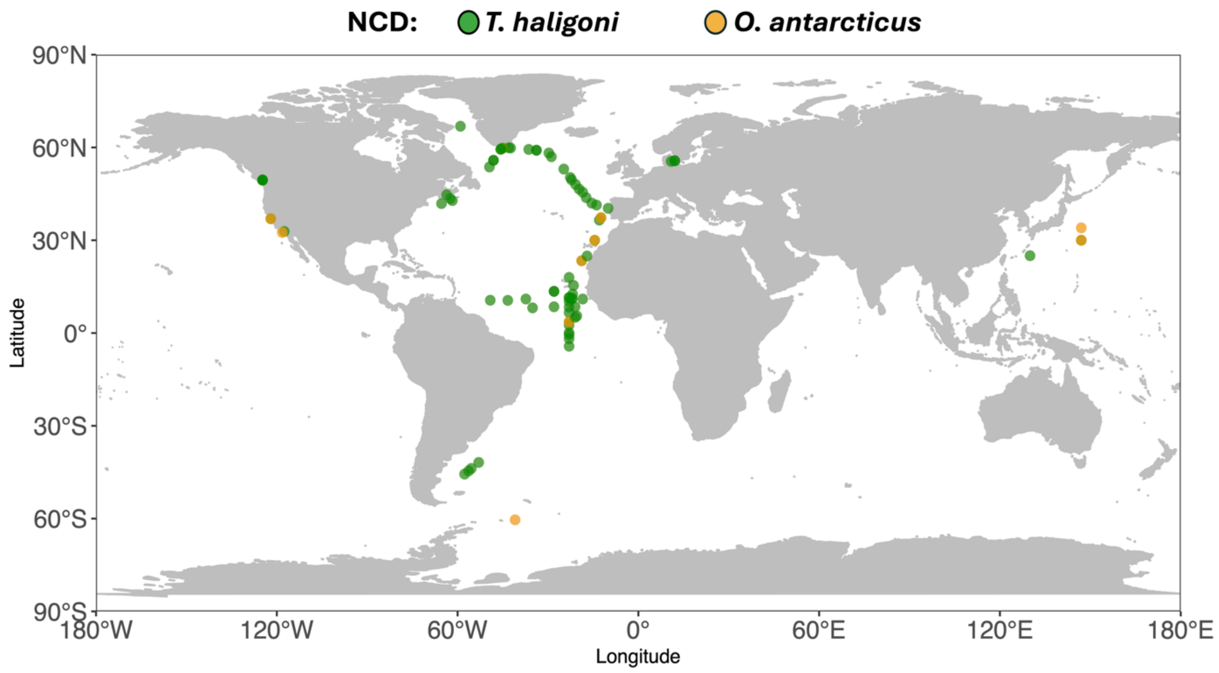


**fig S7** Global distribution *nifH* presence map comparison of *T. haligoni* (green) and *O. antarcticus* (orange) across 865 stations. *nifH* identification for acceptance of species was 100% pairwise identity. The database used for stations can be found at Rose et al., 2024. The figure was modified from Rose et al., 2024 to also include *O. antarcticus* presence and *T. haligoni*.
